# Inclusive pedagogy strategies to introduce high schoolers to systems biology

**DOI:** 10.1101/2023.03.03.531033

**Authors:** Kelsey M Watts, William Richardson

## Abstract

Women and non-white racial and ethnic groups remain underrepresented in science, technology, engineering, and math (STEM). To achieve a more diverse and equitable STEM workforce, the recruitment and retention of these historically marginalized communities in postsecondary education will also need to increase. Recently, the lens has turned to pedagogy and how creating a more inclusive classroom environment can foster STEM identity in previously marginalized communities. This work focuses on developing and piloting systems biology education modules designed to promote an inclusive learning environment in a summer outreach program for high school students by (1) utilizing a hybrid of unplugged activities with coded simulation and (2) a female-oriented problem statement. Based on initial findings in our pilot program, we anticipate that these techniques will enable students with limited prior computational experience to feel more comfortable and able to complete tasks, thereby increasing their self-efficacy and STEM identity. These modules could be a valuable tool for practitioners teaching high school or early college-level computational courses who instruct students with varied coding experience. Additionally, our analysis of the use of a female-oriented problem statement on female-identifying students’ self-efficacy provides potential evidence of the usefulness of representative problem statements in engaging underrepresented student populations. We believe these techniques could be adapted to address a variety of contexts across STEM disciplines and other fields to foster inclusive learning environments.

## Introduction

Women and non-white racial and ethnic groups remain underrepresented in science, technology, engineering, and math (STEM). On average, women make up roughly 50% of the population and hold only a fraction of the engineering and computer workforce jobs (15% and 25%, respectively) [1].

Likewise, historically marginalized communities, including Blacks/African Americans, Hispanics/Latinos, and American Indians, collectively comprise about 31% of the U.S. workforce but only 20% of STEM jobs [1]. Not only are these gender and racial/ethnic gaps leading to inequitable designs, but they are also limiting the ability of the U.S. to compete in the global science and engineering market [2]. Having more women and non-white racial and ethnic groups enter the workforce would allow the U.S. to keep pace with the growing number of STEM jobs and lead to more productive and creative teams that create equitable technology [3, 4].

To achieve a more diverse STEM workforce, the recruitment and retention of these historically marginalized communities in postsecondary education will also need to increase. Although women outnumber men on college campuses, they are less likely to major and graduate with STEM degrees [5]. Likewise, Black and Latinx students who have declared a STEM major are more likely to change majors or drop out than their White peers [6]. These trends are often attributed to the formation of STEM identity, which can be influenced by numerous factors, including middle and high school preparation, societal expectations and stereotypes, university, and workplace culture, representative role models, and STEM self-efficacy [7, 8]. Many informal education and mentoring programs have been created to target underrepresented individuals in an attempt to counteract negative influences and increase STEM identity. However, trends have remained relatively stagnant or even dropped over recent years, indicating that more research needs to be done to prioritize the recruitment and retention of these individuals in STEM [9].

Recently, the lens has turned to pedagogy and how creating a more inclusive classroom environment can foster STEM identity in previously marginalized communities [10]. It is well established that the majority of students benefit from a more active learning environment than traditional lectures [11, 12]. These techniques allow students to learn by doing rather than passively listening. Literature has shown that in a classroom that utilizes active learning techniques, students are less likely to fail the course, which can lead to higher retention rates [13, 14]. However, even in settings that utilize some active learning techniques, such as flipped classrooms and problem-based learning, there can still be additional barriers that impact student performance. For example, previous coding experience in computational classes can predict perceived self-efficacy throughout the course [15, 16]. This trend disproportionately affects those from under-resourced schools and others (like women and non-white racial/ethnic groups) who are less likely to partake in such activities, even if they were an option. These issues have been further exacerbated by the COVID-19 pandemic, which has widened previously existing educational gaps [17].

To address the growing need for computational skills in the workforce and higher education, computational thinking and literacy have become part of the core curriculum for primary and secondary education in recent years [18]. Because the majority of K-12 teachers do not have a lot of previous coding experience, researchers and practitioners have been investigating the use of unplugged activities to facilitate the teaching of computational skills [19, 20]. Unplugged activities facilitate the learning of computational thinking skills through hands-on or role-playing activities in lieu of technology or coding [20–22]. This technique has been effective in increasing primary school students’ computational literacy. Still, little research has been done on how this technique can also benefit older students with limited prior coding experience to increase their self-efficacy in computational methods [22]. Using unplugged activities in conjunction with traditional coding in computational settings could help level the playing field for previously marginalized communities.

Additionally, it is well established that representative role models can increase STEM identity/efficacy for students from historically marginalized populations [23, 24]. However, little research has been done on how representative problem statements (i.e., gender or race-specific) in problem-based learning could also be incorporated into curriculum to increase the self-efficacy and identity of historically marginalized groups. Framing problems as gender or race-specific to target underrepresented student populations may provide them with additional background knowledge, agency, and interest that could influence their STEM self-efficacy and identity formation.

This work focuses on developing and piloting systems biology education modules designed to promote an inclusive learning environment in a summer outreach program for high school students by (1) utilizing a hybrid of unplugged activities with coded simulation and (2) a female-oriented problem statement. We specifically sought to test the following questions:

1. What effect does pairing unplugged learning activities with traditional coding have on the STEM self-efficacy of high school students?
2. To what extent does a female-centered computational biology problem statement impact female high school students’ STEM self-efficacy?

## Methods

### Module Development

We created five systems biology education modules focused on various physiological processes and diseases for high school students. All modules included three major components:

- **Unplugged Activities:** To advance computational thinking and ease students and teachers into modeling, the lesson plans start with a short active learning activity to provide a hands-on or role-playing approach to the phenomena being modeled (i.e., a modified card game to model uncontrolled tumor growth). Each lesson plan also includes recommended discussion questions to connect the unplugged activity with computational concepts.
- **Computational Model Tutorial:** Following the unplugged activity, students work through a guided tutorial to model the biological or disease phenomena. This tutorial leverages the open-source software NetLogo (see Figure 1) [25]. NetLogo was chosen as the platform for model development because of its use of drag-and-drop coding paired with traditional script coding, which is ideal for a target audience of high school students with limited previous coding experience.
- **Computational Model Advancement:** The last component of the modules is a deliverable in which students create their own virtual model by modifying the model they made in the tutorial. The lesson plan provides guided questions as suggestions for model improvement (e.g., What if the patient took a drug that inhibited X, Y, or Z?). It encourages students to develop questions. This last component allows educators to assess gains in their students’ computational abilities.

Each of the five lesson plans created can be found in Appendix A. In addition to the tutorial and model advancement questions given to students, each lesson plan contains an overview page for instructors with learning objectives, definitions of applicable terms, a time to complete estimation, and a guide to the unplugged activity with suggested follow-up discussion questions. All modules are designed to be completed in 2-4 hours each. The unplugged activities are intended to be completed in a group setting. The coding part of each module could be completed individually or as a team. Because they are designed for students with minimal coding experience, the modules build off each other. They are developed to be completed in sequence except for Modules 4 and 5, which cover very similar computational concepts. Students only completed either Module 4 or Module 5 in our pilot study, but in theory, they could complete both if they wanted extra practice on independent module development.

**Figure 1.**
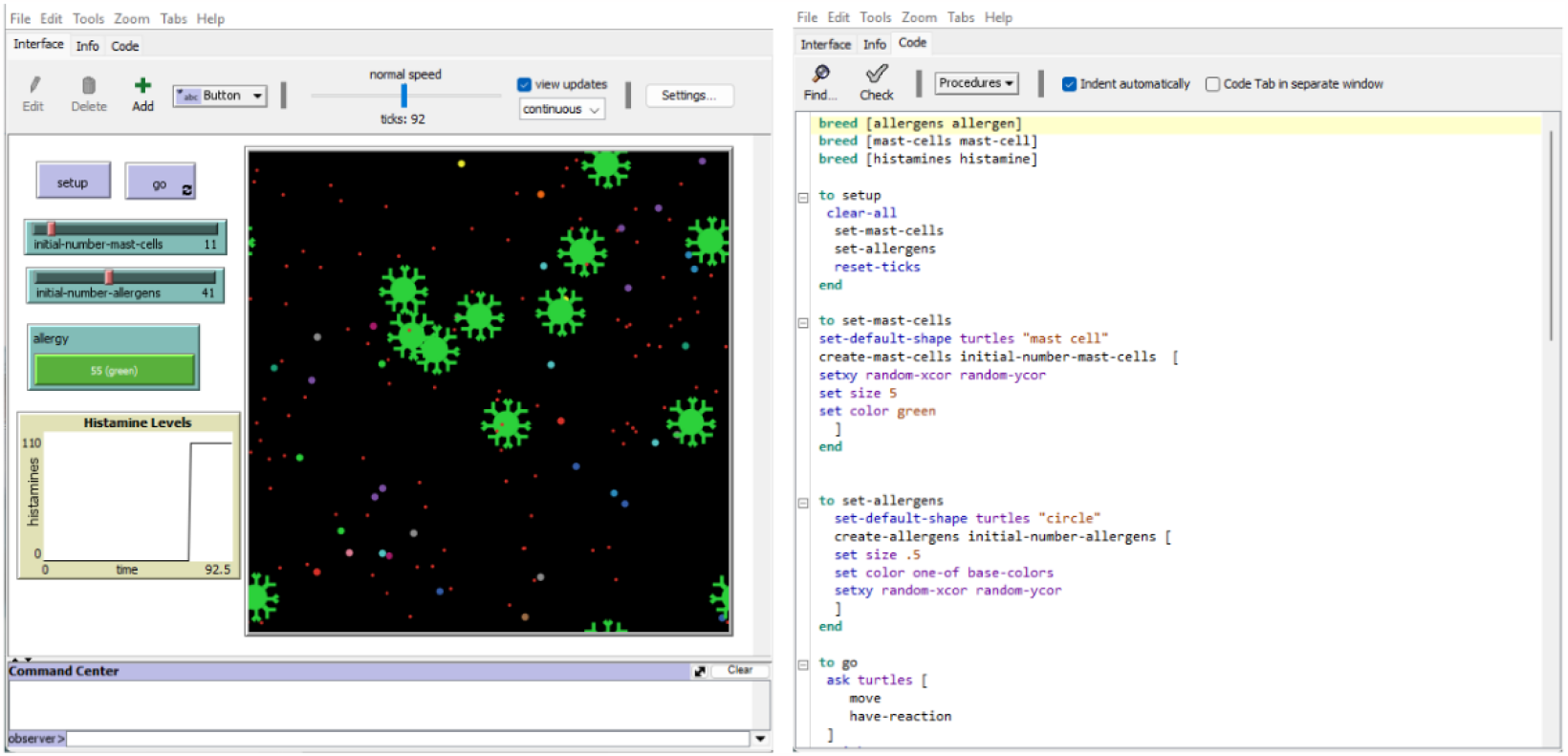
NetLogo interfaces with drag-and-drop buttons and coded script.

Completed NetLogo models can be found on the SysMechBioLab GitHub (https://github.com/SysMechBioLab).

### Modules Created

All of the modules created are centered around common biological phenomena [26]. The first two modules build upon NetLogo’s built-in Tumor and Virus models to ease students into the computational software [27, 28]. Modules 3-5 were independently created and required students to develop a model from scratch (with a guided tutorial). The five modules designed are detailed below and in Figure 2:

- **Module 1: Tumor Growth:** This module employs a built-in NetLogo ‘Tumor’ model to get students acclimated with the environment before asking them to code in it [27]. It simulates a growing tumor and intervention techniques to stop the growth. Biologically, students will observe the progression and treatment of a tumor from a cellular level. Computationally, students will gain experience using a model to test predictions and identify model limitations. The unplugged activity tasks students with playing a modified game of UNO, which simulates the model’s behavior by showing how even if you are down to one card (i.e., a cancer cell), there is still the possibility of regrowth. The model tutorial guides students through interacting with the NetLogo interface. In the model advancement, students are asked to answer questions about the model’s limitations and make suggestions for improvements.
- **Module 2: Virus Prevention:** This module modifies the built-in NetLogo’ Virus’ model to introduce additional parameters that model an intervention to slow the spread of a virus within a population [28]. Biologically, students will examine population dynamics and viral spread. Computationally, students will research and test the effect of input parameters on a model. The unplugged activity guides students through a role-playing game that uses logic comparable to the probability functions which drive the simulation. In the model tutorial, students are shown how to introduce a new variable to the model, ‘mask compliance.’ In the model advancement, students are asked to search the literature to find potential input parameters to model the spread of a virus of their choosing.
- **Module 3: Immune Reaction:** This module guides students through a tutorial to code a NetLogo model from scratch that simulates an allergic reaction. Biologically, students will consider how different cells and molecules interact to elicit an immune response. Computationally, students will define rules and make assumptions to create an agent-based model. The unplugged activity asked students to design the rules for their own game, similar to how they will define rules for the simulation. The model tutorial guides students in setting up the input parameters and all the agents needed for the model (e.g., mast cells, allergens, and histamines). However, the tutorial does not end with a completed model; the agents do not interact with each other. In the model advancement, students are tasked with defining rules for their model and coding them to finish the simulation.
- **Module 4: Gene Regulation:** This module instructs students through a tutorial to code a NetLogo model from scratch that uses Boolean Logic to create a simulation of the lac operon genetic regulation system. Biologically, students will be exposed to a standard model of gene regulation. Computationally, students will define rules using Boolean Logic and analyze model outputs to identify a system’s emergent phenomena. The unplugged activity leads students through a game of If and If Else Simon Says to familiarize them with Boolean Logic [29]. The model tutorial guides students through the beginning steps of creating a model with many interactive parts from scratch. As this simulation is more challenging and extensive than the other modules, students worked in teams during the model advancement for the pilot program to finish the simulation and identify the model’s emergent phenomena.
- **Module 5: The Menstrual Cycle:** This module guides students through a tutorial to code a NetLogo model from scratch that uses Boolean Logic to create a simulation of the menstrual cycle. Biologically, students will be exposed to the hormones involved in the menstrual cycle. Computationally, students will define rules using Boolean Logic and analyze model outputs to identify a system’s emergent phenomena. This module was designed intentionally to mirror Module 4 and was used to investigate the use of a female-centered problem statement vs. a generic problem statement on female-identifying students’ self-efficacy and team dynamics. The unplugged activity, model tutorial, and model advancement utilize similar concepts, and the end simulations are guided by similar logic.

**Figure 2.**
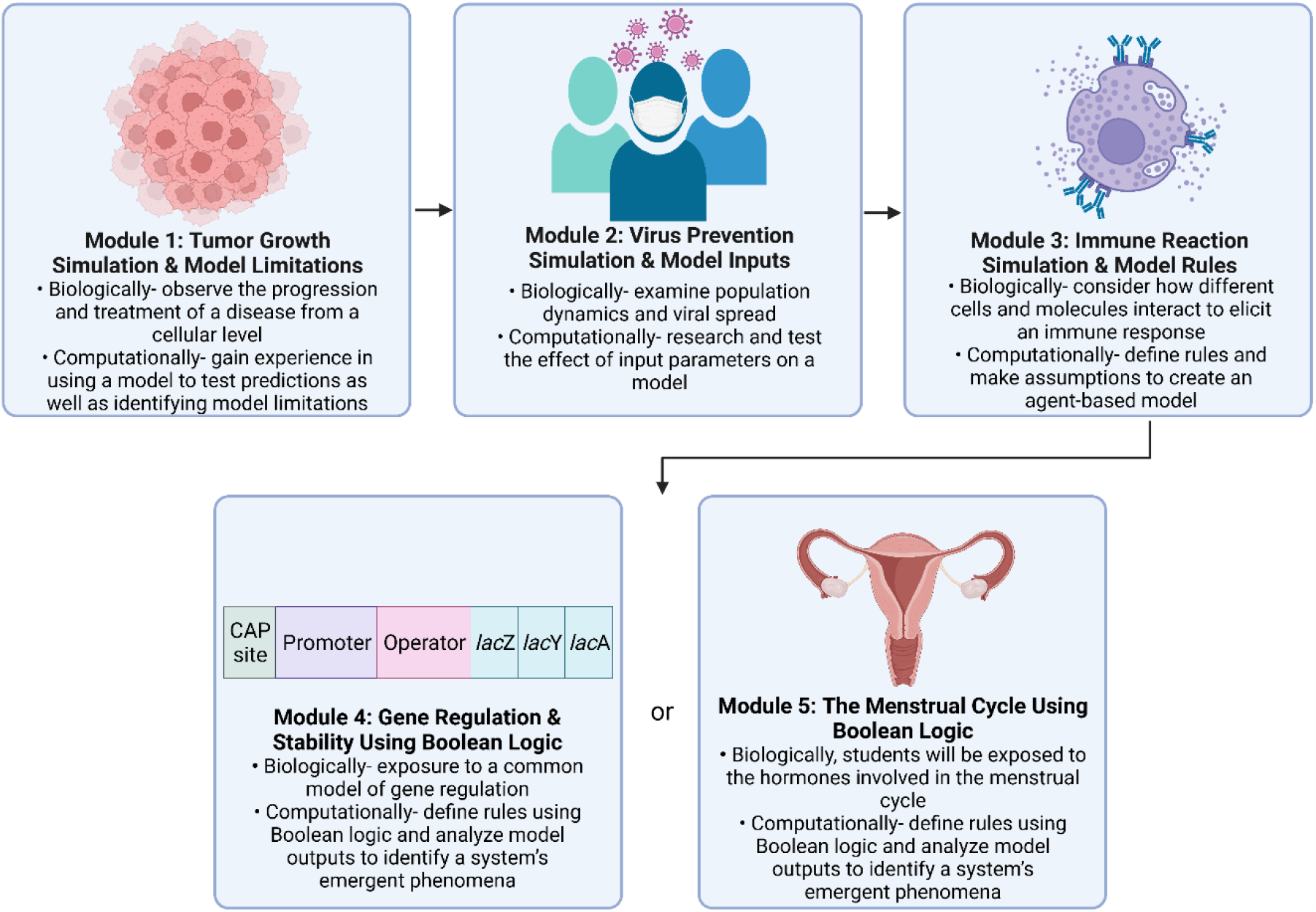
Systems biology modules designed for the program. All students completed Modules 1-3 in sequence and then completed either Module 4 or 5 in the pilot program.

### Pilot Program

The modules were piloted with the Summer 2022 cohort of Clemson’s Emerging Scholars Program. This program invites rising high school sophomores-seniors from some of South Carolina’s most under-resourced schools to campus for several weeks to aid in college preparation. The program helps students prepare for their college applications and has them take classes to expose them to different skills to help them identify potential majors they are interested in. Students enrolled in the program participated in seven 1.5 hr sessions where they completed the modules. All research was conducted in accordance with Clemson’s IRB office and Protocol IRB2022-0059.

The students accepted in the program are most likely to be female or African American/Black, both of which are currently underrepresented in STEM. This provides an ideal sample to study how the inclusive learning techniques incorporated in the modules impact STEM self-efficacy and identity development. In total, 23 students participated in the program. All students self-reported as African American/Black or Mixed race. Thirteen students said they self-identified as female, and five students self-identified as male (4 students did not fill out the survey).

### Quasi-Experimental Intervention

A quasi-experimental approach was used to measure the influence of the female-oriented problem on students assigned to groups to either complete Module 4 (Lac Operon) or Module 5 (Menstrual Cycle) [30]. All students were tasked with completing Modules 1-3 individually. For the final module, students were split up into teams of 4-5, with at least two female students per team. Half of the teams were asked to complete Module 4; the other half were given Module 5.

### Exit Survey and Interviews

An exit survey was deployed on the last day of the program to all students. This survey asked students to rate their pre-and post-understanding of computational and biological learning objectives from the modules using a 1-5 Likert scale [31]. Additionally, the exit survey asked students to rate their confidence and satisfaction with various module components, including the unplugged module activities and the female-oriented problem. Open-ended questions allowed students to provide more depth to their answers. Results from the exit survey were used to (1) evaluate students’ overall satisfaction with the modules, (2) determine if students believed the unplugged activities affected their perceived self-efficacy in completing computational tasks, and (3) determine if a female-oriented problem affected students’ perceived self-efficacy in completing computational tasks. The exit survey can be found in Appendix B.

### Data Analysis

Data were cleaned and analyzed using Microsoft Excel. Mean Pre and Post ratings ± one standard deviation were calculated for each learning objective. A one-tailed unequal variance t-test (not all students answered each question, so a paired test could not be used) between each Pre/Post learning objective was conducted with α=0.1. Mean confidence ratings were also determined for four groups: (1) Female-identifying students who completed the lac operon module (n=5); (2) Female-identifying students who completed the menstrual cycle module (n=8); (3) Male identifying students who completed the lac operon module (n=3); and (4) Male identifying students who completed the menstrual cycle module (n=2). Open-response qualitative data were analyzed for themes and relevant quotes.

### Positionality Statement

The researchers acknowledge the influence that their identities and experiences can impact their studies, perceptions, and reporting of results; therefore, we would like to disclose our positionality [32]. The first author identifies as a white ciswoman early career biomedical engineering researcher. As a woman, she recognizes firsthand how gender can impact prior experience and confidence in completing computational tasks in a male-dominated field. The last author is a white cisgender man in a mid-career engineering faculty position with ∼6 years of teaching experience in computational modeling and systems biology. Together, the research team is devoted to creating more inclusive learning environments for students as a way to strive for equity in STEM education.

## Results

### The modules increased students’ understanding of systems biology concepts

The modules were piloted with Clemson’s Emerging Scholars Program in Summer 2022. The course took place in 1.5 hr sessions over 7 days. In total, 23 students were enrolled in the program and consented to participate in the study. On the program’s final day, an exit survey was administered to assess student gains in understanding biological and computational concepts and perceptions of the modules and the unplugged activities and representative problem statements. The survey had a 78% response rate, with n=18 out of 23 students submitting the survey.

First, gains in understanding were measured by comparing Pre/Post understanding ratings of the learning objectives of the modules. Table 1 outlines the average Pre and Post student ratings for the various biological and computational concepts. On average, every learning objective received a higher understanding rating between the Pre and the Post, with 11/15 of these ratings being statistically significant upon conducting a t-test with α=0.1. Before participating in the class, ratings fell between Minimal Understanding and *Confused Understanding*, but after completing the course, all ratings ranged from *Confused Understanding* to *Good Understanding*.

**Table 1:** Mean ratings ± standard deviation of students’ Pre/Post understanding of various biological and computational elements (1= *No Understanding*, 2= M*inimal Understanding*, 3= *Confused Understanding*, 4= *Good Understanding*, 5= *Excellent Understanding*). A one-tailed unequal variance t-test between each Pre/Post learning objective was conducted (*p*≤0.1*, *p*≤0.01**).

| Learning Objective | Pre | Post | *p*-value |
| --- | --- | --- | --- |
| 1. Describe basic cancer cell metabolism and migration | 2.8 ± 1.4 | 3.7 ± 0.8 | 0.05* |
| 2. Utilize NetLogo's built-in sample models | 2.8 ± 1.5 | 3.5 ± 1.0 | 0.11 |
| 3. Develop and test predictions of therapeutic targets using a tumor model | 2.4 ± 1.3 | 4.0 ± 1.1 | 0.01** |
| 4. Evaluate a model's limitations | 3.0 ± .14 | 4.0 ± 0.6 | 0.01** |
| 5. Describe the factors that can impact a virus's spread throughout a population | 2.8 ± 1.2 | 3.8 ± 1.2 | 0.05* |
| 6. Modify a given NetLogo model by adding and deleting buttons from the main interface | 3.1 ± 1.5 | 3.2 ± 1.2 | 0.43 |
| 7. Research the literature to supply model with input parameters from published experimental data | 2.5 ± 1.3 | 3.5 ± 1.2 | 0.06* |
| 8. Determine what additional rules and/or variables should be added to a model to address its limitations | 2.8 ± 1.3 | 4.0 ± 1.1 | 0.03* |
| 9. Describe the body's immune response due to an allergen | 2.8 ± 1.3 | 4.0 ± 1.1 | 0.03* |
| 10. Define rules for agents of a model given a biological phenomenon | 2.9 ± 1.5 | 4.0 ± 1.1 | 0.04* |
| 11. Code simple rules/commands using NetLogo | 3.0 ± 1.3 | 4.0 ± 1.1 | 0.05* |
| 12. Evaluate how assumptions can increase a model's limitations | 2.8 ± 1.3 | 4.0 ± 1.1 | 0.02* |
| 13. Describe components that make up a biological system and how they function | 3.0 ± 1.3 | 3.7 ± 1.4 | 0.16 |
| 14. Identify stability in a biological process and computational system | 2.8 ± 1.3 | 3.7 ± 1.0 | 0.06* |
| 15. Evaluate model outputs to determine emergent phenomena | 2.6 ± 1.5 | 3.3 ± 1.2 | 0.12 |

### Students valued the inclusive pedagogy strategies

Students were asked if they believed the unplugged activities helped increase their understanding of the modules. Overwhelmingly, 15/18 students said they liked the unplugged activities and cited reasons such as helping them make connections and identify problems more efficiently. Representative quotes include:

> *“Having hands-on activities helped me to understand the biological and computational concepts through a kinesthetic way*.*”*
>
> *“We got to see and understand how everything was supposed to work before we actually built the programs. We could easily identify problems*.*”*

Additionally, the female-identifying students who were assigned the Menstrual Cycle module were asked to what extent did having a female-oriented problem affected their confidence in being able to contribute to the team, and 6/8 students said they thought the menstrual cycle modules increased their confidence, citing prior knowledge made them more quickly understand the problem. Representative quotes include:

> *“Personal experience and prior knowledge helped in completing the task at hand.”*
>
> *“I was familiar with what the hormones did*.*”*

The two male-identifying students who completed the Menstrual Cycle model were also asked if having a female-oriented problem impacted their confidence to contribute to the team, and both said that they did not feel like it affected it at all, one saying:

> *“I gave a good contribution because that’s just the kind of scholar I am*.*”*

After completing the course, students were also asked if they would be more interested in majoring in computational biology/biomedical engineering. Four of the 19 students surveyed expressed that they were more interested in pursuing it as a major.

### Having a female-oriented problem increased female-identifying students’ confidence in completing systems biology tasks

To determine if there was any difference in confidence for students who completed the Lac Operon Module versus the Menstrual Cycle Module, average confidence ratings were determined for male and female-identifying students (Table 2). Of the eight prompts given, on average female-identifying students were more confident in 6 of the areas after completing the Menstrual Cycle module compared to the Lac Operon model. Although no statistical analysis could be conducted because of the small sample size, we believe this indicates a clear trend, especially when paired with the qualitative data. It is important to note, the same trend was not present for male-identifying students, who had a more even split on higher confidence ratings between the Lac Operon and Menstrual Cycle modules.

**Table 2:** Female- and male-identifying students who completed both the Menstrual Cycle and Lac Operon modules were asked to rate their confidence in their ability to carry out various biological and computational components (1= *No Confidence*, 2=*Minimally Confident*, 3=*Moderately Confident*, 4=*Extremely Confident*). The higher ratings within each group are in bold.

| **Computational/Biological Concept** | **Female**<br>**Lac**<br>**Operon** | **Female**<br>**Menstrual**<br>**Cycle** | **Male**<br>**Lac**<br>**Operon** | **Male**<br>**Menstrual**<br>**Cycle** |
| --- | --- | --- | --- | --- |
| 1. Explain computational terms to your peers | 2.2 | **2.6** | 3.0 | **3.3** |
| 2. Explain biological terms to your peers | 2.5 | **2.6** | 2.5 | **3.7** |
| 3. Code a simple biological process using NetLogo individually | 2.3 | **2.6** | 2.5 | **2.7** |
| 4. Code a simple biological process using Netlogo as part of a team | 2.8 | **3.2** | **3.0** | 2.7 |
| 5. Break down complex problems into smaller, simpler problems | 2.5 | **2.6** | **3.0** | 2.3 |
| 6. Make connections between similar problems and experience | 2.5 | **2.8** | **3.5** | 2.3 |
| 7. Identify important information while ignoring unrelated or irrelevant details | 2.8 | 2.8 | **3.0** | 2.7 |
| 8. Design an algorithm to solve a problem | **2.5** | 2.2 | **3.0** | 2.7 |

## Discussion

This manuscript discusses creating and piloting Systems Biology Modules that utilize inclusive pedagogy strategies. We piloted the modules in Summer 2022 with Clemson’s Emerging Scholars Program, an outreach program for high school students. Although we only looked at the effect of the modules on high school students in an outreach program, we believe the modules and inclusive pedagogy strategies could be easily transferable to be used in a typical classroom setting, including an undergraduate entry-level coding/computational biology course. The inclusive pedagogy strategies, in particular, could be of interest to the wider community given the recently proposed changes by Accreditation Board for Engineering and Technology (ABET) to incorporate diversity, equity, and inclusion (DEI) into general engineering programs [33].

The unplugged activities used in the course were well received by students, with 15/18 students saying they enjoyed them and helped them better understand and troubleshoot the computational concepts. Although they may have limited technical relevance depending on the topic, using the unplugged activities was a helpful way to initiate student engagement and buy-in at the beginning of each lesson.

Additionally, the majority of the female-identifying students appreciated the use of a female-oriented problem statement for the final module. It was promising to see their confidence ratings be higher than those with a generic problem statement. However, the female-identifying students’ confidence ratings were still lower than the average male-identifying students’ confidence ratings. This was most apparent in the ratings for explaining computational/biological concepts to your peers, where female-identifying students were *Minimally* to *Moderately Confident*, and male-identifying students were *Moderately* to *Extremely Confident*. This was not entirely surprising, as studies have shown that men typically rate their perceived self-efficacy higher than women in STEM settings [34]. A one-week course is likely not enough to revolutionize the confidence gap in STEM. Future research could investigate the use of representative problem statements over extended periods.

Problem-based learning is often paired with team-based learning in STEM classrooms as a powerful tool to increase motivation and overall learning gains [35, 36]. However, this presents issues for marginalized communities. Stereotypes can be leveraged against them, and task allocation for the technical aspects of a project is not always equal [37]. Based on the initial finding from our pilot study in which a female-oriented problem increased self-efficacy in female-identifying students more than those with a generic problem statement, we believe intentionally designed representative problem statements (i.e., gender or race-specific) could also impact team dynamics such as task allocation, team interactions, team contributions, and having relevant knowledge, skills, and abilities by providing students from underrepresented populations with the additional agency to take the lead. Future studies should not only expand the design of other representative problem statements beyond the one used in this study, but also look into the potential effects on team dynamics and how that may impact self-efficacy.

Limitations of this study include that it was a very small sample size. Due to COVID restrictions, the program was smaller than usual, so we were limited to piloting it with only 23 students. In the future, we hope to continue the modules with more classes/outreach programs to continue testing the influence of inclusive pedagogy strategies. Additionally, our current study only focuses on the immediate effects of the learning modules on student understanding and confidence in completing systems biology tasks. The Emerging Scholars Program does collect longitudinal data on college and major choice, so in the future, we could continue to investigate if the modules have any longer-term effects on students. Also, we only conducted a one-week course as part of the pilot study. Future studies could explore the use of these pedagogies throughout a semester-long course and conduct follow-up studies on major retention and degree attainment to determine these strategies’ long-term effects. Finally, although our student population was 100% Black/African American/Mixed, we did not investigate any possible impact on the intersectionality of race and gender or other factors in our study. This could be an especially interesting future direction regarding the representative problem statements.

## Conclusion

The learning modules developed allow for the teaching of computational techniques through the use of unplugged activities paired with coded simulation. Based on initial findings in our pilot program, we anticipate that this will enable students with limited prior computational experience to feel more comfortable and able to complete tasks, thereby increasing their self-efficacy. These modules could be a valuable tool for practitioners teaching high school or early college-level computational courses who are instructing students with varied coding experience. Additionally, our analysis of the use of a female-oriented problem statement on female self-efficacy provides potential evidence of the efficacy of representative problem statements targeted at underrepresented student populations. We believe such problem statements could be developed to address a variety of contexts across STEM disciplines and other fields to foster an inclusive learning environment.

## Acknowledgments

Thanks to Dr. Lisa Benson for providing guidance on the development of the modules and experimental protocols. All figures were created with BioRedner.com.

## Notes

https://github.com/SysMechBioLab

### Competing Interest Statement

The authors have declared no competing interest.

